# Factors Affecting Detection Of GMO Traces In Food And Crop Samples

**DOI:** 10.1101/869032

**Authors:** Waqar Hassan, Nadia Jamil, Shehzaib Siddiqui, Anam Ali, Owais Quadri, Maliha Wajeeh, Aftab Ahmed Khan, Saifullah Khan

## Abstract

Rice is one of the highly consumable and demanded food crops throughout the world. To meet this requirement, genetically modified (GM) crops were introduced, which were met with sturdy and frequently angry opposition by the user. To handle the situation labeling of GM food / crop was advised, thus demand for traceability and detection of unknown modified genes increased.

Modified genes are detected by various methods; these methods are affected by several factors as reported in international standards. Those factors include presence / absence of shells, husk, and dust, method of DNA extraction and different PCR primers. These factors ultimately pose hindrance in detection of modified genes.

This study was conducted to analyze the effects of the above mentioned factors on detection of GM rice samples. Furthermore two different sets of primers were used with same samples and their impact on the PCR detection was observed.

The results showed a significant difference in DNA concentration between the rice samples with husk, without husk and after seed germination. Furthermore, the change of PCR primer set also affected the detection of genetic modifications. This allows analysis of the potential factors that may have impact on the final results.

## I. Introduction

Genetic modifications in food crops are trending due to the rising demand for resistance from pests, increase in yield, and fortification of crop (Bouis et al, 2003). Globally, manipulation of food crops at genetic level and production of GM crops is under debate and no consensus has been attained upon their human consumption due to unforeseeable consequences regarding multiple ethical, legal and health issues (Cook & Robbins, 2004; Pretty et al, 2010). Furthermore the list of approved GMOs varies from country to country and the control of unapproved GMOs is necessary to screen the occurrence of GMOs that are not authorized (Cankar et al. 2008). These issues were discussed in convention on biological diversity (CBD), and the Cartagena protocol was adopted. This protocol aims to ensure the safe handling, transport and use of living modified organisms (LMOs) that may or may not have adverse effects on biological diversity, also risks to human health were taken into account (Gabol et al., 2012). The Cartagena protocol forces the governments to notify in advance in case of trade of GM crops. Protocol allows banning the unsafe GM products and places a stipulation of labeling shipments that can cause any threat to traditional crops and environment (David, 2001).

European Union and Asian countries unlike USA have always had their qualms on the use of GMOs (Gabol et al., 2012) and due to the serious concerns among the consumers, cultivation of GM crops and their application in food products for human consumption is strictly prohibited (Kardum et al, 2017). In the past, transformation of rice seeds for pest resistance *via* genetic modifications were practiced without proper labeling, as a consequence GM Rice was reported by many international markets, and presence of cry1/2 genes was reported (De Steur et al, 2017; Park, Kim & Kim, 2017). Due to rise in unauthorized genetically modified genes (UGMs) it was felt mandatory to properly label food item as GM and Non-GM (Boccaletti et al, 2017), and certify all crop imports as GM free, particularly in European Union, China, and Pakistan (Clement et al, 2017; Faour & Todd, 2018).

Recently few labs have submitted their work on GMOs affecting human health and may be in near future the exact effect of GMO foods on human or animal health can be established (Lee et al., 2017).

Pakistan has 0% tolerance policy for GM food. The routine genetic modifications that are monitored avidly includes, screening for presence of Cauliflower mosaic Virus 35S gene (CaMV 35S) commonly known as promoter. Another gene that is commonly screened alone or along with CaMV 35S is Nopaline Synthase (NOS) terminator gene from *Agrobacterium tumifaciens* also known as tNOS or terminator, terminator region of cauliflower mosaic virus is also used but less common. ISO 21569:2005 documents the PCR detection of CaMV 35S (promoter) and *Agrobacterium tumifaciens* NOS (terminator) as standard for GM crop screening. Therefore this method was adopted by CESAT as well.

However after discussion with other labs in vicinity it was felt that the screening based on promoter and terminator region can be affected by the use of different primers and same samples can be detected as GM on the basis of one set and non-GM on the basis of other primer set. These factors are other than the already documented factors like processing of samples, methods of DNA extraction, limit of detection of PCR, and presence of PCR inhibitors in samples (Bonfini et al, 2001) to name a few.

Keeping aforementioned in view, a study was designed on rice samples and effect of following factors was observed on DNA concentration and screening of GM crop; 1) extraction of DNA from rice with husk (WH) and without husk who) and germinated seedlings (GS), 2) effect of DNA extraction method on DNA conc. and 3) effect of using two different primer sets for same samples.

## II. RESULTS

DNA was extracted from 11 rice seed samples WH, WoH and GS (Table 1). Results showed that there was significant difference between extracted DNA conc. of samples (Table 1).

**Table 1.** DNA Conc. of Samples

| S.No | Sample ID | With Husk |  | Without Husk |  | After Germination |  |
| --- | --- | --- | --- | --- | --- | --- | --- |
|  |  | Commercial KIT ng/μl | CTAB Method ng/μl | Commercial KIT ng/μl | CTAB Method ng/μl | Commercial KIT ng/μl | CTAB Method ng/μl |
| 1 | 48 | 33.820 | 16.250 | 34.610 | 69.950 | 72.060 | 14.050 |
| 2 | 55 | 21.180 | 18.950 | 26.080 | 77.700 | 45.930 | 3.200 |
| 3 | 80 | 14.900 | 20.650 | 59.610 | 30.950 | 118.300 | 9.700 |
| 4 | 81 | 86.180 | 31.450 | 0.196 | 20.800 | 88.240 | 4.250 |
| 5 | 82 | 94.800 | 26.750 | 34.020 | 55.300 | 32.840 | 5.800 |
| 6 | 248 | 33.820 | 14.900 | 29.610 | 81.350 | 52.650 | 3.600 |
| 7 | 250 | 11.570 | 19.050 | 20.290 | 36.500 | 32.060 | 14.400 |
| 8 | 296 | 58.730 | 41.500 | 38.430 | 36.550 | 73.820 | 43.050 |
| 9 | 342 | 97.550 | 25.000 | 19.410 | 35.050 | 108.800 | 4.950 |
| 10 | 351 | 125.900 | 20.800 | 24.800 | 30.050 | 62.750 | 2.650 |

Keeping in view aforementioned samples were used for screening the effect of two different primer sets. To make best use of the system, most commonly used transgenic elements were selected as PCR target i.e. cauliflower mosaic virus (CaMV) 35S promoter and *Agrobacterium tumefaciens* Nopaline synthase terminator (tNOSs).

The primer sequences are mentioned in Table 2. All samples were positive for internal positive control. Primer pair 35S-1/-2 (promoter) and nosFMZP1/2 (terminator) showed the bands at 195 and 125 bp respectively (Fig. 2), while primer pair 35SF/R (promoter) and HA-nos118-f/r (terminator) failed to show any positive bands at expected 269 and 118 bp (Fig. 3).

**Table 2.**
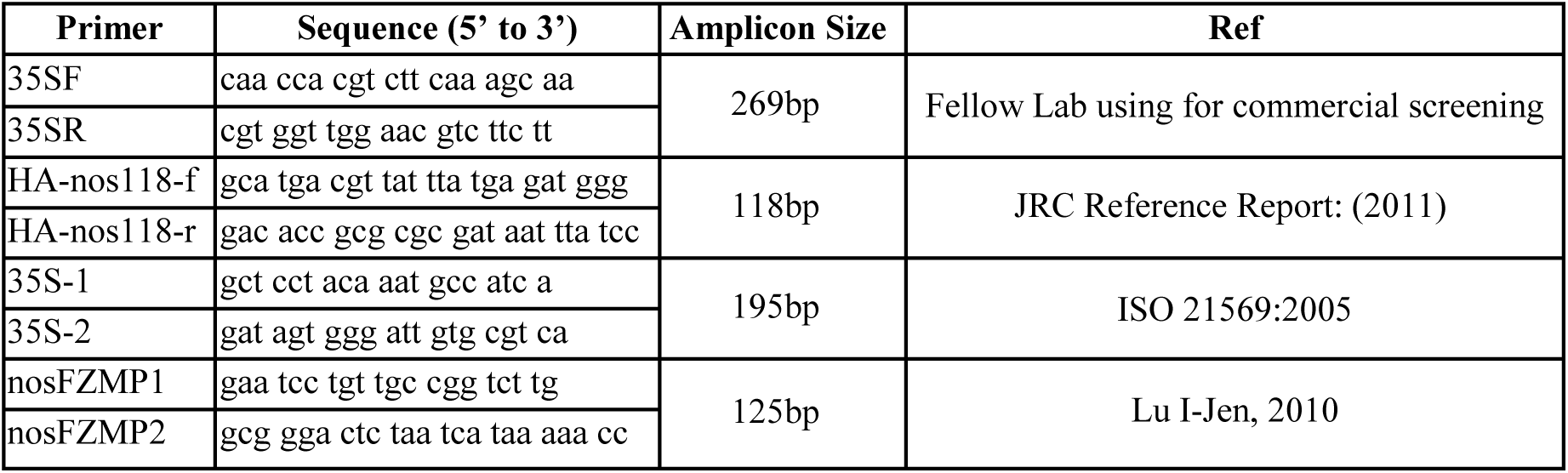
Primers used in this study

**Fig. 1:**
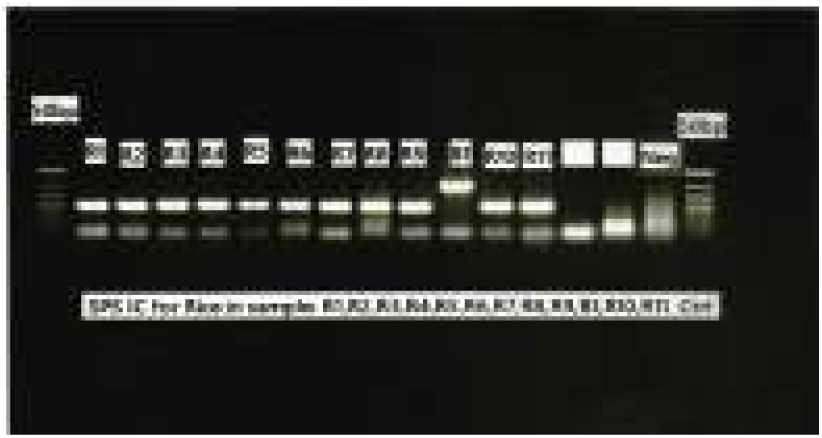
Gene amplification of IC of Rice Key: R1-R9: Rice samples, B1: Known GMO Cotton sample, R10-R11: Rice Samples

**Fig. 2:**
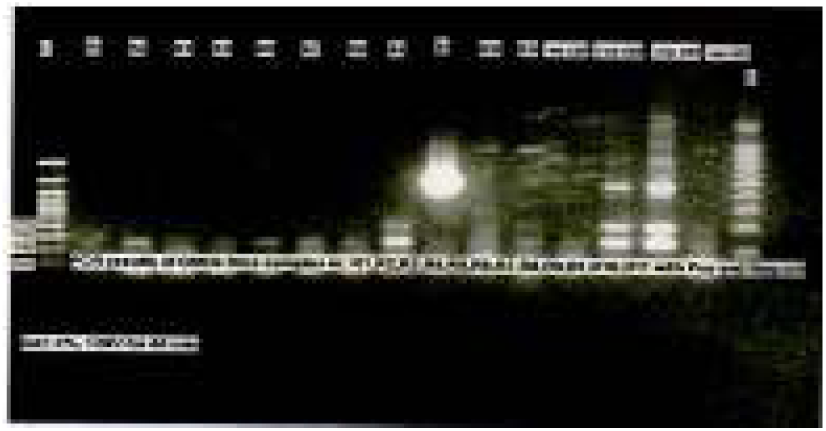
Duplex PCR for Gene amplification of promoter and Terminator region in Rice Key: R1-R9: Rice samples B1: Known GMO Cotton sample R10-R11: Rice Samples

**Fig. 3:**
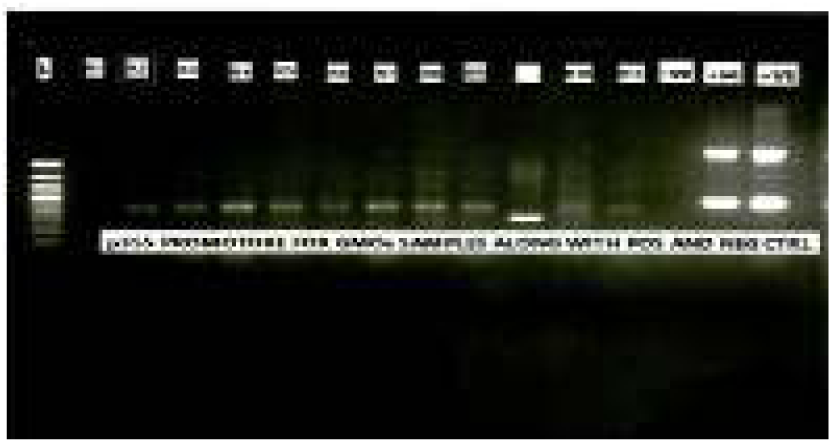
Duplex PCR for gene amplification of promoter region in Rice Key: R1-R9: Rice samples B1: Known GMO Cotton sample R10-R11: Rice Samples

## III. DISCUSSION

To overcome the challenges arising from the rapid increase in the number of GM crop products, preliminary screening of GM crop is gaining importance day by day. According to information from the AGBIOS biotech crop database, approximately 139 GM crops in total have been commercially approved throughout the world. These crops are modified for as much as 20 genes, thus increasing the number of possible modified genes per crop (James, 2015).

Many processing steps affect the state of the DNA conc. however in many cases, PCR amplification will be achievable. Although, assessment of effect by different factors on DNA concentration seems easy but comparison of results is not.

The best method of detection of genetically modified genes is PCR and aptness of isolated genomic DNA as study material for PCR based detection of GMOs depends upon the quality of the DNA. However on one hand the choice of a certain extraction method affects the quality or quantity, on other hand the choice of primer may affect the outcome. Therefore care should be taken when results are compared. The number of cycles, the length of amplicon, and DNA amount supplemented in a PCR reaction also affect detection and quantification limits. Short amplicons of 150 bp maximum should be given preference and whenever available, internationally authenticated PCR assays should be used (Gryson, 2010). During this study effect of presence or absence of husk and seedling germination on DNA concentration of 11 rice seed samples were studied by DNA extraction with commercial kit and CTAB method described in ISO standard.

The results suggest that DNA concentration increases by both methods if the samples are used WOH. However according to Romer Labs, washing of GM samples or removing the shell or dust from sample is not allowed and may produce incorrect results (Futrnyk, 2017).

Keeping in view, aforementioned WH samples were used for screening the effect of two different primer sets for promoter and terminator region on same samples.

All PCR reactions were performed in triplicate and duplex PCR detection system was used to improve the efficiency of screening process.

Primer pair 35S-1/-2 and nosFMZP1/2 showed the bands of promoter and terminator at 195bp and 125bp respectively, while primer pair 35SF/R and HA-nos118-f/r failed to show any positive bands at 269bp and 118 bp. Few unexpected PCR products appeared when duplex PCR method was applied, that could be due to some transgene construct having same sequence stretch as the target one where primer got attached and amplification occurred (Fraiture et al, 2015). There is also one more thing to be noticed that the GM region is modified in a way that the primer did not recognize the site and got attach at another region therefore in-line with suggestions of JRC scientific and technical reports a method / primer with low specificity can have a higher probability of detection with any chosen method (EUR 25008 EN-2011).

## IV. Conclusion

The increase in number and divergence of GMOs developed and commercialized has gradually forced the GM detection laboratories to rationalize their analytical work and rise of UGMs is demanding that one method of screening is not enough for giving ultimate decision about presence or absence of genetically modified genes.

## V. MATERILAS AND METHODS

### DNA Extraction

11 rice seed samples were taken each sample was in duplicate (with husk and without husk). 150mg of sample was taken and homogenized in separate sterile tubes to fine powder. Genomic DNA extraction was performed by CTAB method as defined in ISO-21570 and commercial kit.

### DNA purity and degeneration state

Concentration of genomic DNA was first measure using the “*Multiscan Go*” UV-vis spectrophotometer (Thermo Scientific, USA). Concentration (ng/μl) and A260/A280 readings were recorded for each sample. Quality of the extracted DNA was further verified by electrophoresis on 1% (w/v) agarose gel (TAE/TBE buffer system). The results were visualized on gel documentation system (Azure bio systems, USA).

### Qualitative PCR

Extracted genomic DNA samples were tested for CaMV 35s Promoter and Nos Terminator region by using two sets of primers (Table 1). To amplify target sequences, newly synthesized primers were diluted (10 μmol/L) for further use in PCR. Following reaction mix was made in 25 µl reaction tubes, 3 µl of extracted DNA, 12.5 µl Master Mix (Affymetrix, USA) and 0.5 µl reverse and forward primers of each set were used.

The reactions were performed on Eppendorf Master Cycler Pro (Eppendorf, Germany). Amplification profile was as follows; initial denaturation was done at 95°C for 5 min, followed by 38 cycles of 95°C for 30 sec, 60.5°C for 40 sec, 72°C for 40 sec and final extension at 72°C for 5 min.

### Analysis of DNA Amplified Product

The electrophoresis of the amplified DNA products was carried out in 2% agarose gel containing ethidium bromide 2.5 µl per 50 ml. Gel was subjected to constant voltage of 5 volts/cm in TAE buffer. The gel analysis was performed on gel documentation system (Azure bio systems, USA).

## Acknowledgement

The authors thankfully acknowledge whole CSD group especially Misbah Khadim and Abdul Mutalib Murwat for providing assistance in sample receiving and preparation.

## Author Contribution

WH and NJ gave idea, interpreted results and wrote the paper

MW and SS did the PCR work

AA and OQ did all the DNA extraction work AAK and SK reviewed and corrected the paper

